# Linear ubiquitin chains remodel the proteome and influence the levels of hundreds of regulators in *Drosophila*

**DOI:** 10.1101/2024.05.09.593206

**Authors:** Oluwademilade Nuga, Kristin Richardson, Nikhil Patel, Xusheng Wang, Vishwajeeth Pagala, Anna Stephan, Junmin Peng, Fabio Demontis, Sokol V. Todi

## Abstract

Ubiquitin controls many cellular processes via its post-translational conjugation onto substrates. Its use is highly variable due to its ability to form poly-ubiquitin with various topologies. Among them, linear chains have emerged as important regulators of immune responses and protein degradation. Previous studies in *Drosophila melanogaster* found that expression of linear poly-ubiquitin that cannot be dismantled into single moieties leads to their own ubiquitination and degradation or, alternatively, to their conjugation onto proteins. However, it remains largely unknown which proteins are sensitive to linear poly-ubiquitin. To address this question, here we expanded the toolkit to modulate linear chains and conducted ultra-deep coverage proteomics from flies that express non-cleavable, linear chains comprising 2, 4, or 6 moieties. We found that these chains regulate shared and distinct cellular processes in *Drosophila* by impacting hundreds of proteins. Our results provide key insight into the proteome subsets and cellular pathways that are influenced by linear poly-ubiquitin with distinct lengths and suggest that the ubiquitin system is exceedingly pliable.

## INTRODUCTION

Myriad cellular processes and pathways in eukaryotic cells depend on the small modifier protein, ubiquitin (UB)(1). UB is tethered chemically to other proteins post-translationally. In essence, UB functions by changing the interaction landscape, protein folding, or both properties of its substrates. Because UB can be conjugated to other UB moieties, it can create various types of chain lengths, shapes, and sizes, including branched chains. A tremendous body of work conducted in vitro, in cellular systems, and in vivo has cataloged shared and distinct outcomes of the conjugation of different types of poly-UB to proteins, and has also evidenced critical roles for unanchored “free” ubiquitin chains that are linked head-to-toe, also known as linear UB chains(1, 25).

UB conjugation and de-conjugation are controlled by hundreds of enzymes. The coordinated action of a main E1 (UB-activating enzyme), of E2s (UB-conjugating enzymes) and E3s (UB ligases; figure 1A) ensures the chemical preparation and targeting of the 8.6 kDa UB protein to a specific amino acid on the substrate, most commonly a lysine (K) residue(1). Errors in UB conjugation are resolved by deubiquitinases (DUBs), which are also tasked with UB recycling for maintaining a pool of free UB(12). The overall notion is that UB is conjugated in a stepwise manner onto a preceding UB to generate chains of different lengths and shapes. UB removal is conducted either one-by-one or *en bloc*, the latter then being dismantled to single UB proteins, ensuring a UB pool for reutilization (12, 14, 26).

**Figure 1:**
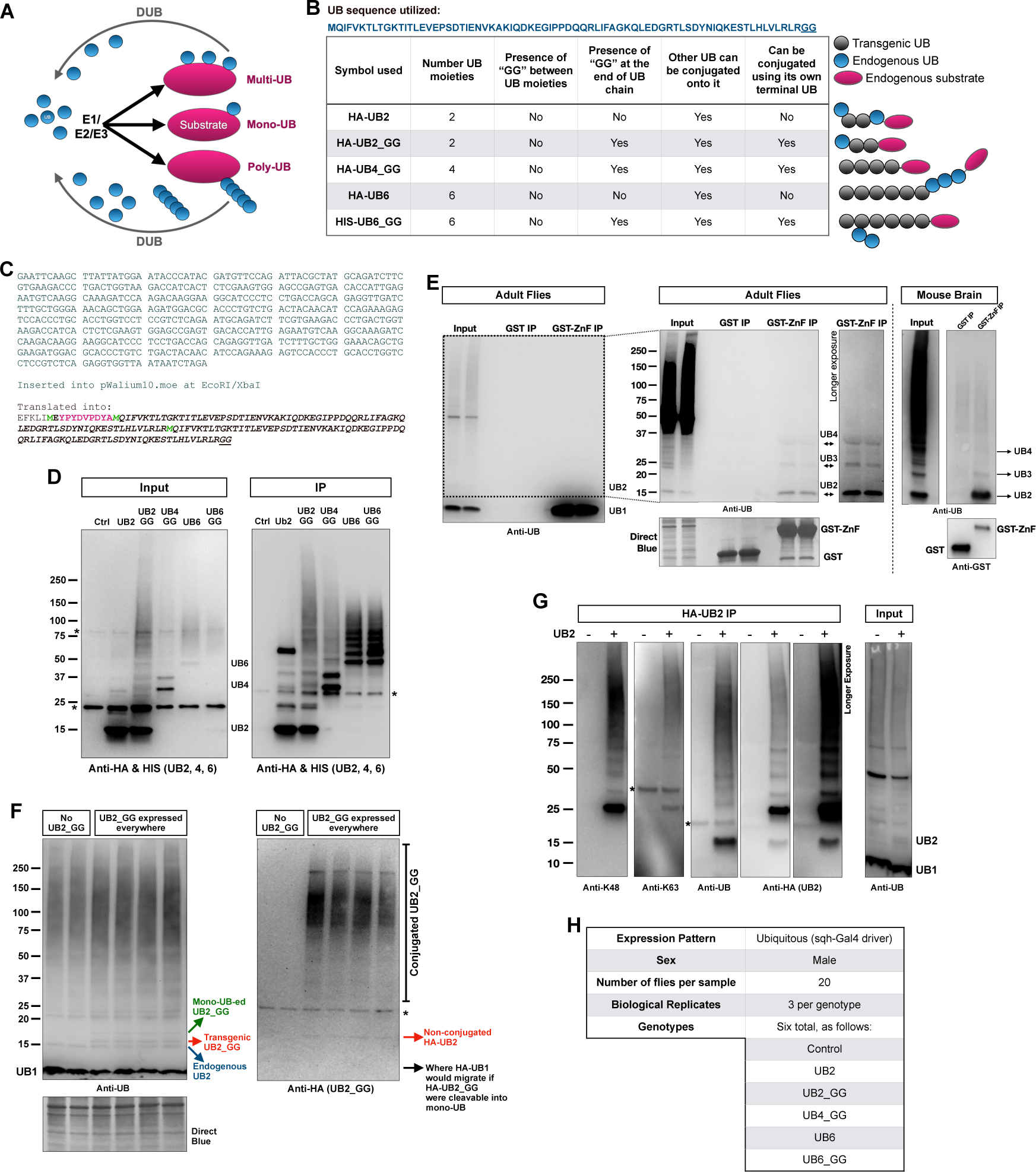
A) Graphical summary of the ubiquitin conjugation and deconjugation system. B) Tabulation of the different linear UB chains that were over-expressed in flies and their properties. Legend and graphics on the right portion of this panel indicate types of conjugation that may occur with the transgenic linear UB species. C) Nucleotide and amino acid sequence of the UB2_GG construct. The other lines generated for this project were designed similarly. D) Western blots from the expression of the linear UB chains used in the study. The lysates were from male flies of the same crosses used for proteomics. Immunoprecipitation procedures are in the methods section. “Ctrl” line was the sqh-Gal4 driver in trans with the host line used to generate the UBX transgenics. E) Isolation of unanchored ubiquitin using recombinant GST-tagged ZnF domain from USP5, processed as outlined in the methods section. In Western blots from flies, the vast majority of unanchored UB was mono-UB. Cutting the membrane to remove this species (dotted box) and exposing it longer revealed other unanchored UB species. For blots from mouse brain, mono-UB was cut off from the membrane to reveal the other chains. Mouse brain sample was leftover lysate from mice used in our prior publication (68). The input and IP lanes from the mouse brain lysate blots are from the same membrane and exposure, cropped and rearranged to ease visualization. F) Western blots from male flies expressing the noted UB chain. G) Immunoprecipitation and Western blots from flies expressing UB2_GG everywhere. H) Summary of the groups used for proteomics. In Western blots, asterisks denote non-specific bands.

To expand the general understanding of the ways in which UB can function *in vivo*, we generated a series of transgenic *Drosophila melanogaster* UB chain-expressing lines that comprise 2, 4 or 6 UB in a head-to-toe arrangement. The fly lines are all isogenic to each other, with each transgene inserted into the attP40 site on chromosome 2 to ensure equal expression (20, 27-29). These transgenic fly lines express distinct versions of linear poly-UB of different lengths: one type lacks a terminal “GG” that would enable its conjugation onto another protein, whereas the other type contains a terminal “GG” motif and is able to be conjugated onto other proteins *in vivo* (27). Our prior studies indicated that expression of linear poly-UB is not detrimental to *Drosophila*, regardless of the expression pattern; that these chains are degraded by the proteasome; that chains with a terminal “GG” can be conjugated onto other proteins; and that chains without a terminal “GG” are decorated by numerous types of linkages comprising endogenous UB (20, 27-29). Collectively, the expression of these exogenous chains suggests a highly flexible and adaptable UB system, able to handle linear UB chains that cannot be removed via the action of DUBs (20). However, a key outstanding question is whether there are specific sets of proteins that are particularly sensitive to modulation by linear UB chains.

Here, we addressed this gap in knowledge by identifying the cellular processes and pathways that are impacted by the expression of distinct linear poly-UB that can or cannot be conjugated onto protein substrates. We hypothesized that some of these UB chains could be utilized to enhance the degradation of specific proteins, and that UB chains that cannot be conjugated onto other proteins could also regulate cellular processes independently from protein degradation. Through the utilization of quantitative, unbiased proteomics (30) of male flies that expressed UB2, UB4 or UB6 chains ubiquitously, we found that linear UB chains distinctly influence cellular processes and molecular functions. Collectively, our results expand the appreciation of the organism’s flexible use of UB and identify proteins that are particularly sensitive to modulation by linear poly-UB.

## RESULTS

### Expression of different linear UB chains in *Drosophila*

We previously reported the generation of transgenic fly lines containing UB6 chains that cannot be dismantled due to the absence of “GG” motifs between constituent moieties(27, 29). These chains (figure 1B; 1C details UB2 and all the other lines were generated similarly) were designed to lack (or contain) a final “GG”, meaning that they could not (or could) themselves become conjugated onto endogenous proteins in the fly through this terminal motif. Here, we expanded the toolkit to examine the function of linear UB chains by generating three additional transgenic fly models containing UB2 linear chains without or with the terminal “GG” and UB4 with a terminal “GG”. All of these lines are isogenic and were site-integrated (with the phiC31 system(31)) into attP40 on *Drosophila* chromosome 2 to ensure equal expression. Each line is expressed through the binary Gal4-UAS system (32) and the sqh-Gal4 driver, which enables transgene expression in all fly tissues, throughout development and in adulthood(33). All analyses were conducted with male flies that were 10 days old. Figure 1D shows the expression of the various UB transgenes in these flies.

We decided to generate transgenes that express UB2, UB4 and UB6 chains for the following reasons. UB4 is generally accepted as the minimal length required to effectively target a substrate for proteasomal degradation(34). Based on biochemical work that we conducted with whole fly tissue or mouse brain lysates, we noticed that among unanchored chains isolated using the Zinc finger domain of USP5, which binds unanchored UB(6, 35), UB2 was the most prominent species (figure 1E; please note that UB1 was vastly more abundant than unanchored UB2 in this assay). We also reasoned that it would be informative to examine the effect of the next largest building block of UB, after UB1, in the overall proteome. As shown in figures 1F and 1G, UB2_GG is tethered with other proteins in the fly and is used to generate mixed-linkage chains, indicating its incorporation into the UB pool *in vivo*. Lastly, we thought it reasonable to also test UB6 in this study, both as a logical next, longer UB chain, and because we had tested its impact in the fly before(27, 29). With these tools on hand, we proceeded to conduct unbiased, quantitative proteomic analyses.

### TMT mass spectrometry of fruit flies expressing non-cleavable linear UB chains

We conducted 18-plex tandem mass tags (TMT) mass spectrometry from flies expressing the above-mentioned linear UB chains ubiquitously during development and in adulthood. We used the same driver we have used in other studies with these chains, sqh-Gal4. The groups that we used for examination were: UB2, UB2_GG, UB4_GG, UB6, and UB6_GG (figure 1H). The control group with no UB chain expression consisted of flies with the sqh-Gal4 driver on the same genetic background used to generate the linear UB transgenics.

Groups were tested in triplicate per genotype, with 20 flies per group. Approximately 100 µg protein per sample were used per group. The TMT tags used in each group are shown in supplemental figure 1. We acquired 1,843,638 MS/MS spectra, identified 74,120 unique peptides corresponding to 7,907 unique proteins with a false discovery rate of <1%, and quantified 7,872 unique proteins with at least 2 spectral counts per protein. The Principal Component Analysis plot derived from these data is shown in figure 2A, providing a bird’s-eye-view of similarities and differences among the assessed groups and replicates.

**Figure 2:**
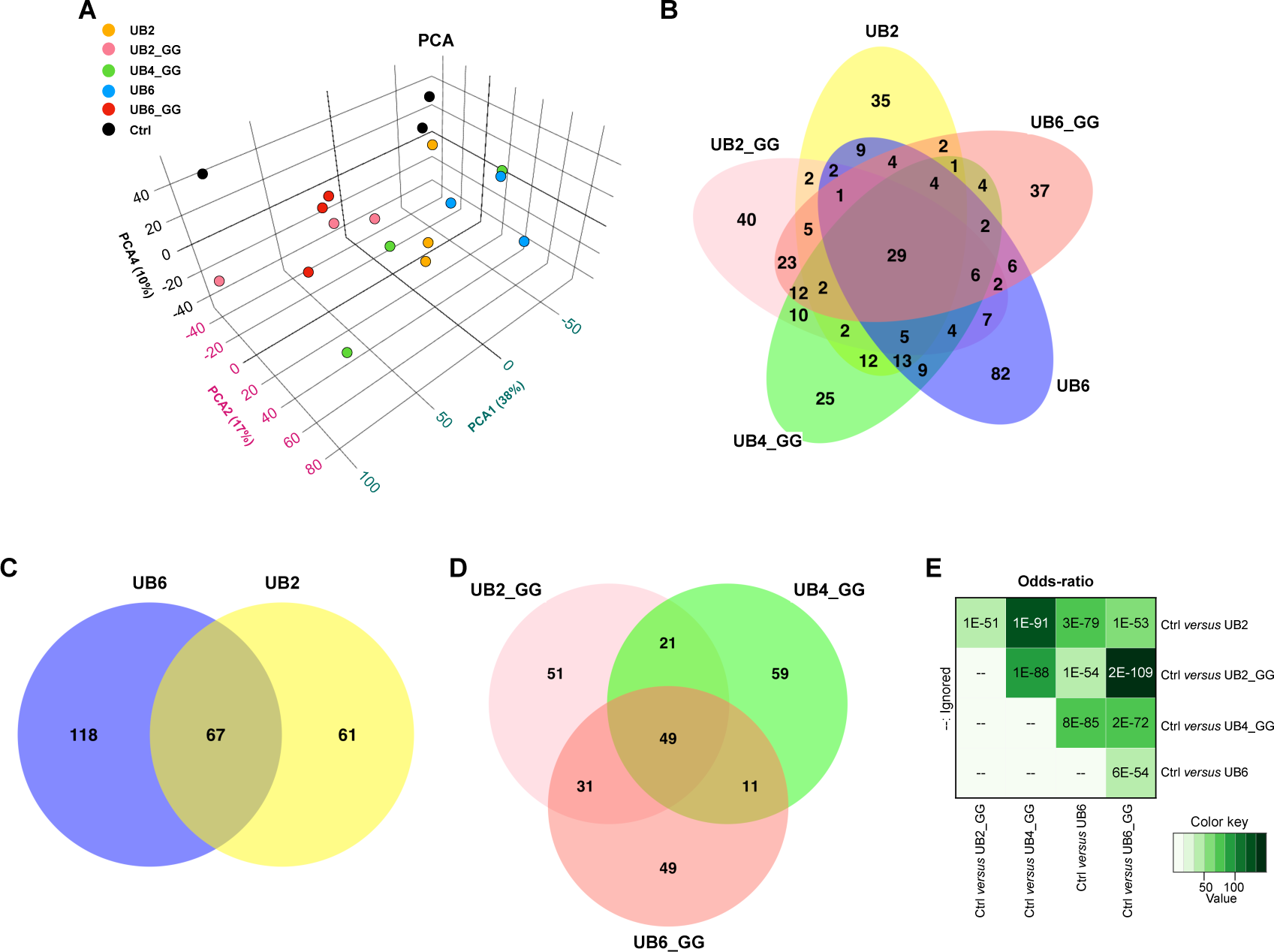
Principal Component Analysis (A), Venn diagrams (B-D) and protein overlap with odds ratio analyses (E) of the differentially abundant proteins relative to control group observed between and among the different UBX groups, collectively indicating shared and distinct proteins among the groups.

First, we compared all linear UB-expressing groups together to determine the extent to which the differentially abundant proteins in each group overlap with other constructs; we observed significant similarities among them (figure 2B, 2C, 2D). There are more extensive similarities between groups that have conjugation potential (UBX_GG) than length of ubiquitin chain (UB2 *versus* UB2GG, or UB6 *versus* UB6GG) (figure 2E).

Twenty-nine proteins were shared among all groups (supplemental figure 2 lists their identities), compared to the control group (figure 2B). There was a unidirectional decrease in 13 proteins and increase in 16 proteins across all groups (figure 2B and supplemental figure 2). These included CYP4p1, CYP4ac, GstE5 and Ugt36E1, which are involved in detoxification and hormone metabolism (36).

Each linear UB-expressing group also had between 25-82 proteins whose changes in abundance were unique to them (figure 2B). We further sub-categorized these groups into linear UB without or with a terminal “GG” motif. When comparing UB2 with UB6 lacking a terminal “GG”, there were approximately equal numbers (67 shared proteins; figure 2C and supplemental figure 3), including a decrease in proteins involved in fatty acid metabolism, e.g., ACBP3 and ACBP5, which enable fatty Acyl-CoA free pool of long chain fatty acid accumulation in the cytosol (36) as well as a decrease in enoyl-CoA hydratase (CG4594), which is involved in the breakdown of fatty acids via beta oxidation (36). This was accompanied by a decrease in proteins predicted to enable ecdysteroid 22-kinase activity, as well as LSP2 which is inducible by ecdysteroids(37) and an increase in chitin and chitin interactors (CG13155 & CG32240). Furthermore, there was an upregulation of CG34717 (acyl-CoA reductase in *Drosophila*), which is predicted to function in long-chain fatty-acyl-CoA metabolic and energetic processes (36). There were also differentially abundant proteins unique to each group (118 for UB6 and 61 for UB2; figure 2C).

When comparing the other linear UB chains that contain a terminal “GG” motif (figure 2D), we observed 49 shared proteins among all groups, including increased abundance of CG11598 (lipase activity) cryptochrome (circadian feedback loop regulator) as well as a 2.5 log_2_FC increase in Ugt36E1 (involved in sex pheromone discrimination when expressed in the olfactory sensory neurons(36)). Conversely, we identified a decrease in the protein levels of alpha-Est9 and CG4594 enoyl CoA hydratase (figure 2D and supplemental figure 4).

The Venn diagrams in figure 2 indicate that there are distinct and shared proteomic changes when each of the transgenes encoding for linear UB chains is expressed ubiquitously in *Drosophila*. Odds-ratio analysis provides a statistical assessment of these overlaps (figure 2E), which indicates highly significant cross-regulation among the linear UB groups when compared to the control line. The degree of significance varies, with the highest overlap being observed between UB2 and UB4_GG, and the least between UB6 and UB6_GG. Collectively, these results demonstrate specific changes in the fly proteome caused by the expression of distinct linear UB chains. Supplemental figure 5 lists all the protein changes, compared to the control flies.

### Molecular and pathway analyses of the TMT proteomics data

Pathway enrichment analyses of differentially abundant proteins highlighted key molecular functions and biological processes that were significantly altered by the expression of each of the linear UB species (figure 3). As shown in figure 3A, expression of each of the linear UB species led to up- or down-regulation of specific molecular functions in the fly. Some such changes (oxidoreductase activity, monooxygenase activity, iron ion binding, heme binding, tetrapyrrole binding, glutathione transferase activity, and transference of aryl and alkyl groups) were shared among 4 out of 5 of the different linear UB types. Expression of linear UB chains led to 4 or more (UB2_GG, UB4_GG, UB6) changes in functions, with UB2 and UB6_GG expression resulting in the largest number of such changes (10 and 9, respectively). Thus, expression of the linear UB chains of different lengths in the fly led to shared and distinct changes in molecular functions. When comparing UB chains without a terminal “GG” *versus* poly-UB with a terminal “GG”, there was considerable overlap in the molecular functions modulated by UB chains, except for 3 pathways (oxidoreductase, long chain fatty acyl-CoA binding, and carbon-nitrogen activity) which are present in UB2 but absent in UB2_GG. UB2_GG in turn was enriched for serine-type peptidase activity while UB2 was not. Overall, UB4_GG displayed the fewest changes in molecular functions (figure 3A).

**Figure 3:**
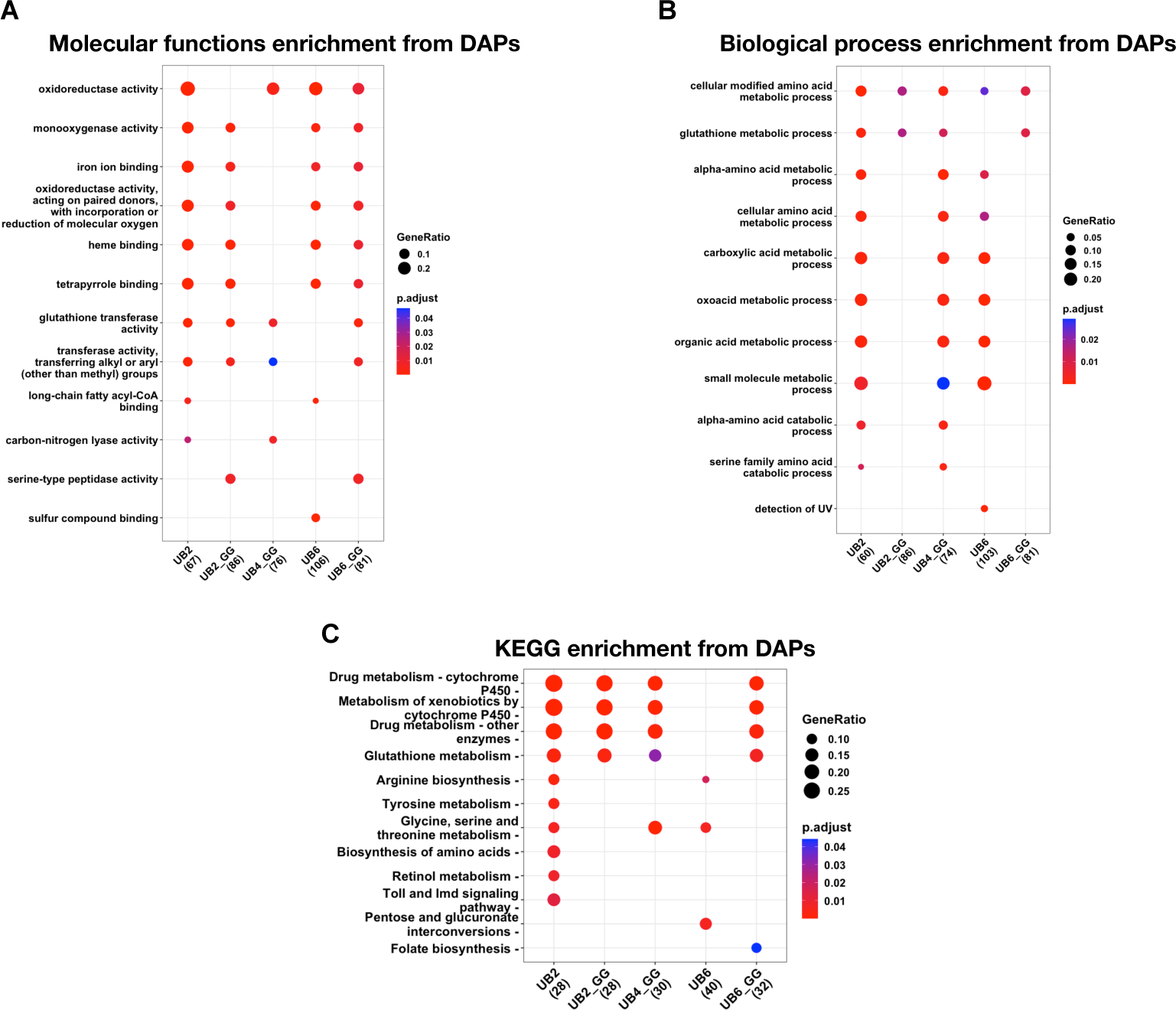
Gene Ontology pathway enrichment analyses of differentially abundant proteins showing over-represented molecular functions (A), biological processes (B), and KEGG pathways (C) changed by the expression of specific linear UB species in flies. Key differences arise among the different groups expressing specific types of linear poly-UB, especially when comparing chains with a “GG” versus not.

Overlaps and differences among the linear UB chains can be further contextualized when examining biological processes (figure 3B). UB2 and UB4_GG led to the largest number of changes in biological processes, a total of 10, mostly centered around energy metabolism, amino acid metabolism, and detoxification. UB2_GG and UB6_GG had the fewest, a total of 2 each, that were shared with UB2, UB4GG and UB6_GG (cellular modified amino acid metabolic process, glutathione metabolic process).

KEGG pathway enrichment also showcased UB2 (figure 3C) as the group with the most changed pathways (10), with the other groups varying from 3 (UB6) to 5 (UB4_GG and UB6_GG). Four pathways were shared among all groups, except for UB6: drug metabolism-cytochrome P450, metabolism of xenobiotics by cytochrome p450, drug metabolism-other enzymes, and glutathione metabolism. Four pathways were exclusive to UB2 (tyrosine metabolism, biosynthesis of amino acids, retinol metabolism, and Toll and IMD signaling pathway). UB6 had one pathway altered only in this group, pentose and glucuronate interconversions, whereas folate biosynthesis was altered only in UB6_GG. Collectively, these results again indicate shared and distinct processes influenced by linear UB of distinct lengths in *Drosophila*.

### Curated assessment of the proteomics data – the ubiquitin/proteasome system

We next took a curated look at processes generally linked to ubiquitin. We began by comparing ‘ubiquitin mediated catabolic processes’ among the 5 linear UB groups (figure 4A). The levels of many proteins were changed, albeit not markedly so overall. We observed that several 20S proteasome core alpha and beta subunits were increased across all linear UB groups, but to different extents depending on the exact component (the 20S core component is the degradative machinery of the proteasome (38)). We also observed changes in 19S proteasome cap subunits, including Rpn and Rpt proteins (the 19S regulatory cap component of the proteasome binds to, deubiquitinates, and unfolds substrates targeted for degradation (38)). In some cases, their levels were generally higher (e.g., Rpt3, Rpt4) and in other cases, they were lower in specific groups (e.g., Rpn9 in UB2_GG, UB6_GG) but higher in others (Ub2, UB6), potentially reflecting specific modifications to the functions of the 19S regulatory cap on a needs-basis, depending on the homeostatic status of the cell. Changes in the levels of various ubiquitin enzymes and ubiquitin-binding proteins were also detected across the different linear UB groups.

**Figure 4:**
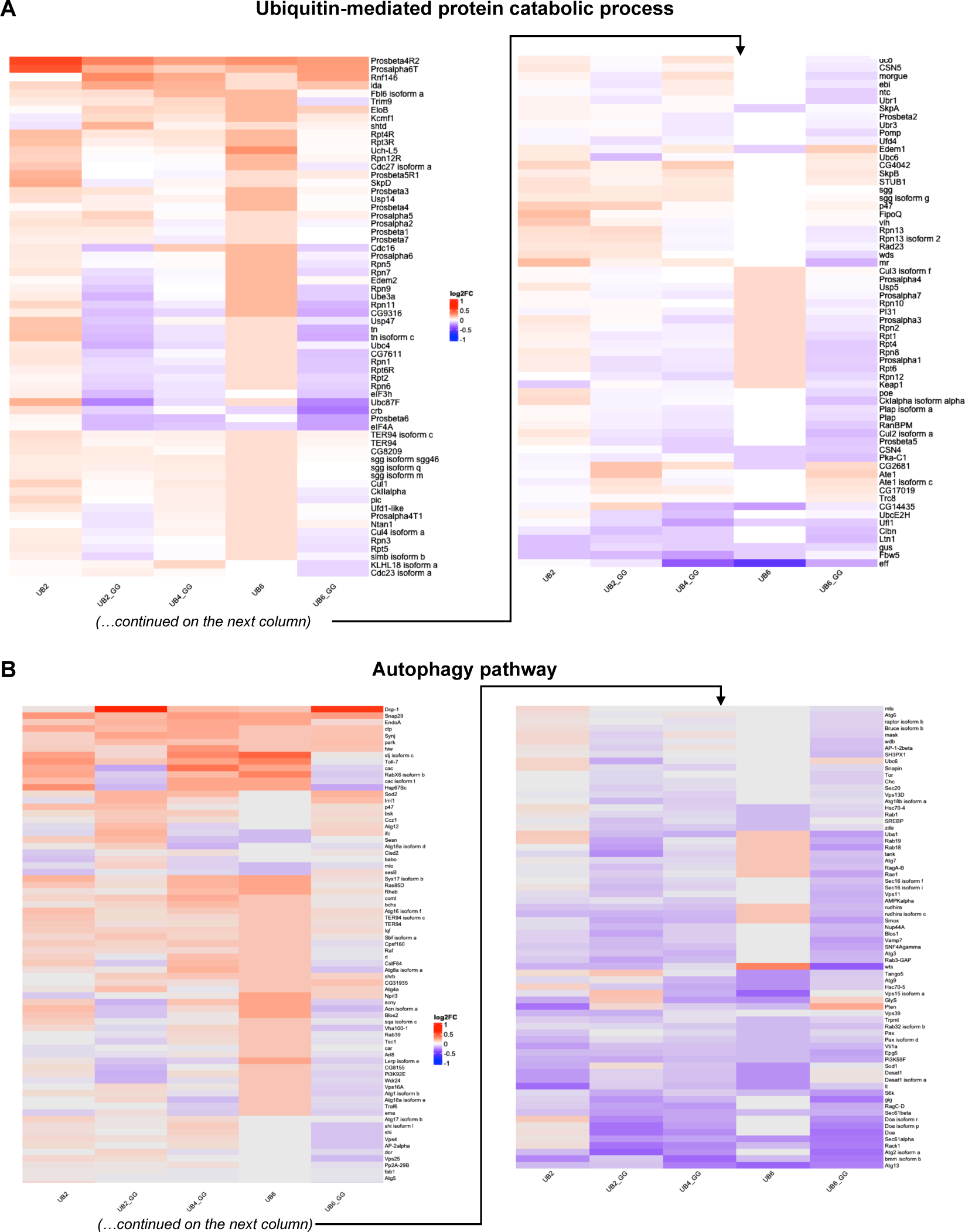
A) Heatmap of hierarchically clustered, TMT-quantified protein expression changed in curated ubiquitin mediated catabolic process (flybase.org) in the presence of specific linear UB species with log2FC expression relative to control group. B) Heatmap of proteins changed in autophagy pathway in the presence of specific linear UB species. The levels of various proteasome-, UB-related enzymes and proteins, and autophagy-related components are impacted across the different genotypes.

There were general trends that matched among UB2_GG, UB4_GG and UB6_GG with swathes of proteins consistently regulated across these groups. These included various Rpn and Rpt proteins whose levels were lower, alpha and beta 20S subunits whose levels were higher, and some UB ligases and deubiquitinases whose level also changed in the same direction within these groups. The combination of increased levels of some ubiquitin-dependent proteins *versus* decreased levels of others could be an indication of the utilization of the UBX_GG linear chains in modulating cellular processes and pathways, presumably by targeting specific protein substrates in a manner dependent on the length of the available linear poly-UB.

There were also similarities between UB2 and UB6, whose pattern differed to some extent compared to the “GG” groups. In this case, most of the changes that reached significance were in the upregulated direction and included various proteasome subunits, UB ligases, deconjugases, and UB-binding proteins. We previously reported the degradation of UB6 in *Drosophila* by the proteasome in a manner dependent on ubiquitin-binding proteins p47 and VCP (known in the fly as TER94) (27). The generally increased levels in proteasome subunits, ubiquitin ligases, and ubiquitin-binding proteins (figure 4A) are congruent with the notion of key roles being played by these proteins in the degradation of linear UB species, and of possible adaptations of the proteasome in response to changes in the abundance of linear UB chains. Altogether, these analyses indicate specific adaptations in the ubiquitin-proteasome system in *Drosophila* because of linear UB chains that can be conjugated onto other proteins *versus* UB chains that cannot be conjugated.

### Curated assessment of hierarchical proteomics data – the autophagy pathway

We next examined the hierarchical clustering of protein changes involved in the autophagy pathway, where ubiquitin signaling is also important and includes the utilization of linear UB chains (20). As with the other pathways, there were similarities and differences among the different linear UB groups (figure 4B). The “GG” groups trended more similarly in their overall patterns, especially UB2_GG and UB6_GG. UB2 and UB6 shared some similarities, especially in the bottom one-third of figure 4B, but not as much in other portions. A few proteins stood out for the changes in their levels among the different groups. From the top, Dcp-1 (Death caspase-1, involved in non-apoptotic processes, including autophagy(36)) was upregulated in the presence of UB2_GG and UB6_GG, with minimal or no upregulation in the other groups. Hsp67Bc (a small heat shock protein that stimulates autophagy(36)) had decreased levels with UB2_GG and UB6_GG, but increased levels with the other groups. Another interesting example was wts (warts, a kinase involved in Hippo signaling and tissue growth(36)), whose levels were increased with UB6, but decreased with UB6_GG. Lastly, the levels of Atg13 (Autophagy-related 13, which controls the initiation of autophagosome formation(36)) were lower among all groups, but more so with the longer UB chains. Other examples with differences in their levels among groups exist, including, Sod1, gig, Vps15, Pten, Atg2, etc. Once again, it appears that linear poly-UB with different lengths and capabilities influences a key pathway, in this case autophagy, in specific ways in the fruit fly. In the case of the “GG” groups, the decrease in protein levels is presumably a sign of the conjugation of these protein substrates to UB2_GG, UB4_GG or UB6_GG, which would target them for degradation by the autophagy-lysosome or ubiquitin-proteasome systems. Conversely, the increased levels of other proteins may reflect their increased stabilization via ubiquitination.

### Curated assessment of hierarchical proteomics data – the immune pathway

Since linear ubiquitin is closely involved in the immune response (20), we next focused on changes in immune pathways, represented in a hierarchical heatmap in figure 5A. While there was overlap in the protein levels and their change in direction among the different linear UB groups (e.g., top quartile for UB2, UB2_GG, UB4_GG, UB6, and UB6), differences in patterns also emerged. For example, in the second quartile of figure 5, UB2_GG and UB6_GG were more similar between each other, and UB6 stood out by itself in directionality of change (generally higher). In the third quartile UB2 was less similar than the others, whereas in the bottom quartile UBX_GG were more similar among each other, and UBX were more similar between themselves.

**Figure 5:**
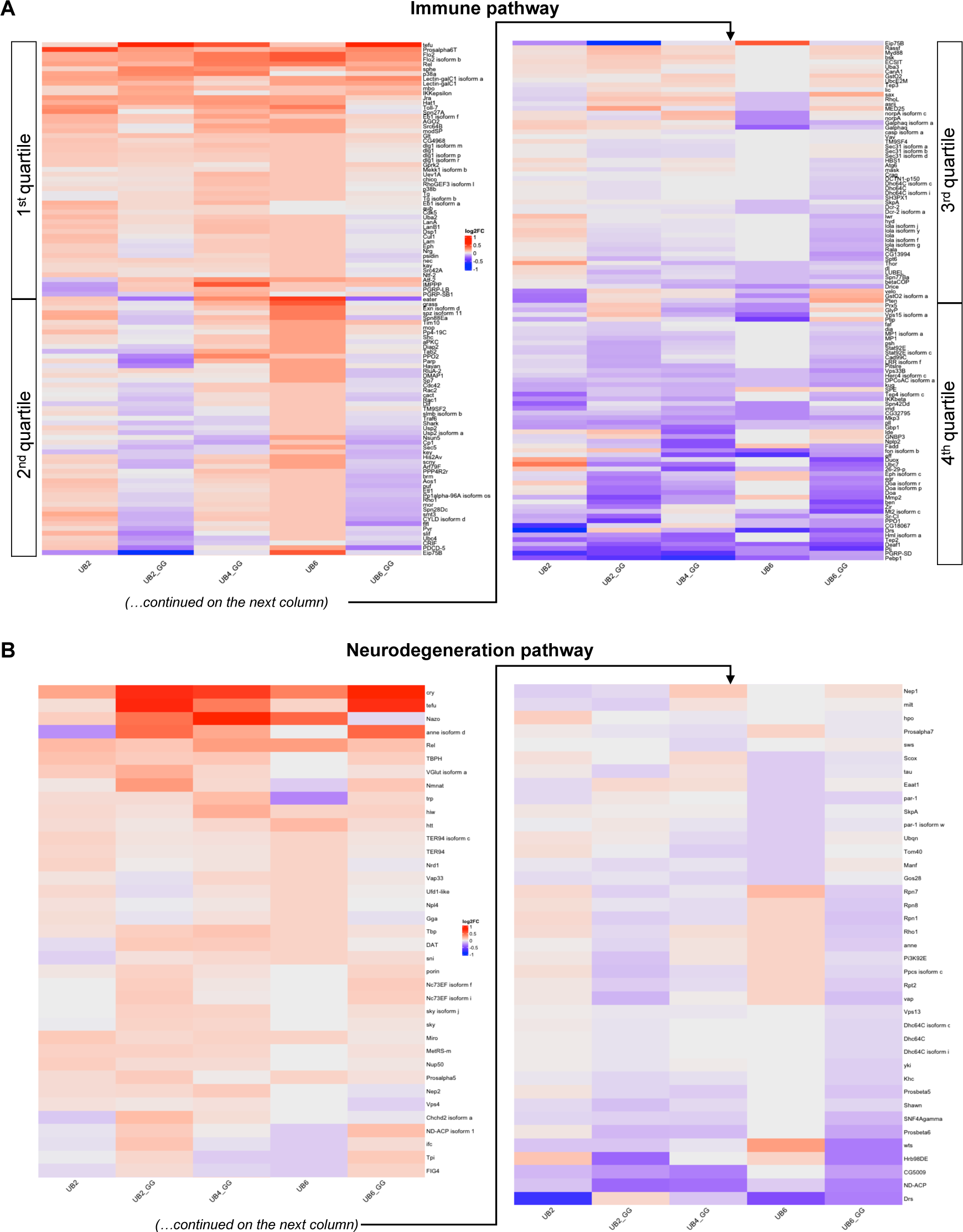
A) Heatmap of hierarchically clustered, TMT-quantified protein expression changed in curated immune pathway (flybase.org) in the presence of specific linear UB chains. Similarities and differences among the groups fall roughly into 4 areas, as highlighted by “quartiles”. B) Heatmap of hierarchically clustered TMT quantified protein expression changed in curated (neurodegeneration pathway) flybase.org in the presence of specific linear UB species. Similarities and differences are evident when comparing linear UB without or with a terminal “GG.

Among the many proteins identified, some stood out in terms of their extent and directionality of change. From the top, one striking example was the protein tefu (telomere fusion, a non-specific serine/threonine kinase(36)), whose levels were upregulated in the presence of UB2_GG, UB6_GG, and less markedly so in UB4_GG, but were not altered as much in the non “GG” groups. Another interesting case was eater (a transmembrane receptor involved in the phagocytosis of Gram-positive bacteria(36)), whose levels were reduced in the presence of UB2_GG and UB6_GG, but increased to varying extents in the other groups. Other similar examples (e.g., PDCD-5, Galphaq, eff, Drs, etc.) collectively suggest distinct outcomes on immune proteins resulting from linear UB chains with different lengths and ability to become conjugated onto other proteins in *Drosophila*.

### Curated assessment of hierarchical proteomics data – the neurodegeneration pathway

Lastly, since ubiquitin-mediated protein quality control is a *bona fide* protective molecular mechanism in neurodegenerative diseases (39, 43), we assessed changes to neurodegeneration pathway proteins upon expression of the UBX and UBX_GG constructs (figure 5B). Although the linear UB constructs were ubiquitously expressed and not targeted to the nervous system, we identified changes in abundance of proteins that have been shown to be neuroprotective in distinct neurodegenerative pathologies, most notably, ‘cry’,‘tefu’, and ‘Nazo’. Cry (cryptochrome; a regulator of the circadian rhythm(36)) was upregulated across all species, but especially in the presence of UBX_GG. Tefu (telomere fusion, a non-specific serine/threonine kinase whose human orthologue, when mutated, causes ataxia telangiectasia(36)) was significantly increased in UB2_GG and UB6_GG. Similarly, Nazo (involved in triglyceride homeostasis(36)) was differentially increased in UB2_GG, UB4_GG, and UB6 with its changes in UB2 and UB6_GG falling below filtering cutoff (supplemental heatmap and raw data spreadsheet). Overall, UB2_GG and UB6_GG share patterns of changes across the board. UB6 stands apart from the other groups, and there are some trends of similarity between UB2 and UB4_GG. Future studies focusing on the nervous system with these specific UB species may reveal insights into the role of linear ubiquitin in brain function, and how chains of specific length may influence the initiation and progression of neurodegeneration.

### Decreased protein levels in flies expressing linear UB with a terminal “GG” motif

As shown in figures 1D, F and G, and as we have published before with UB6_GG (27), expression of linear UB species with a terminal “GG” motif leads to higher molecular weight ubiquitin smears. We do not observe this type of migration with UB2, which lacks a terminal “GG” (figure 1D); we have also shown before that ubiquitin smears of UB6, which also lacks a terminal “GG”, result from conjugation of endogenous UB onto these non-cleavable, non-conjugatable chains (summarized in figure 1B) (27, 29). UB6_GG can be conjugated onto endogenous proteins in the fly and in mammalian cell culture (27). Thus, we reasoned that the higher molecular species that we observe with the “GG”-containing linear UB likely include endogenous proteins onto which UB2_GG, UB4_GG, or UB6_GG was conjugated, a modification that could impact their levels.

Since UB is commonly thought of as a degradative signal, we curated a list of proteins whose levels were consistently decreased among the three groups of linear UB chains with a “GG” motif. As summarized in figure 6, the levels of 26 proteins were significantly reduced in the presence of UB2_GG, UB4_GG, and UB6_GG. Some of them have known functions that span different activities (transcription, cytoskeletal, etc.), whereas a few do not have a known function. Altogether, these findings indicate that linear UB species of different length may share common protein targets.

**Figure 6:**
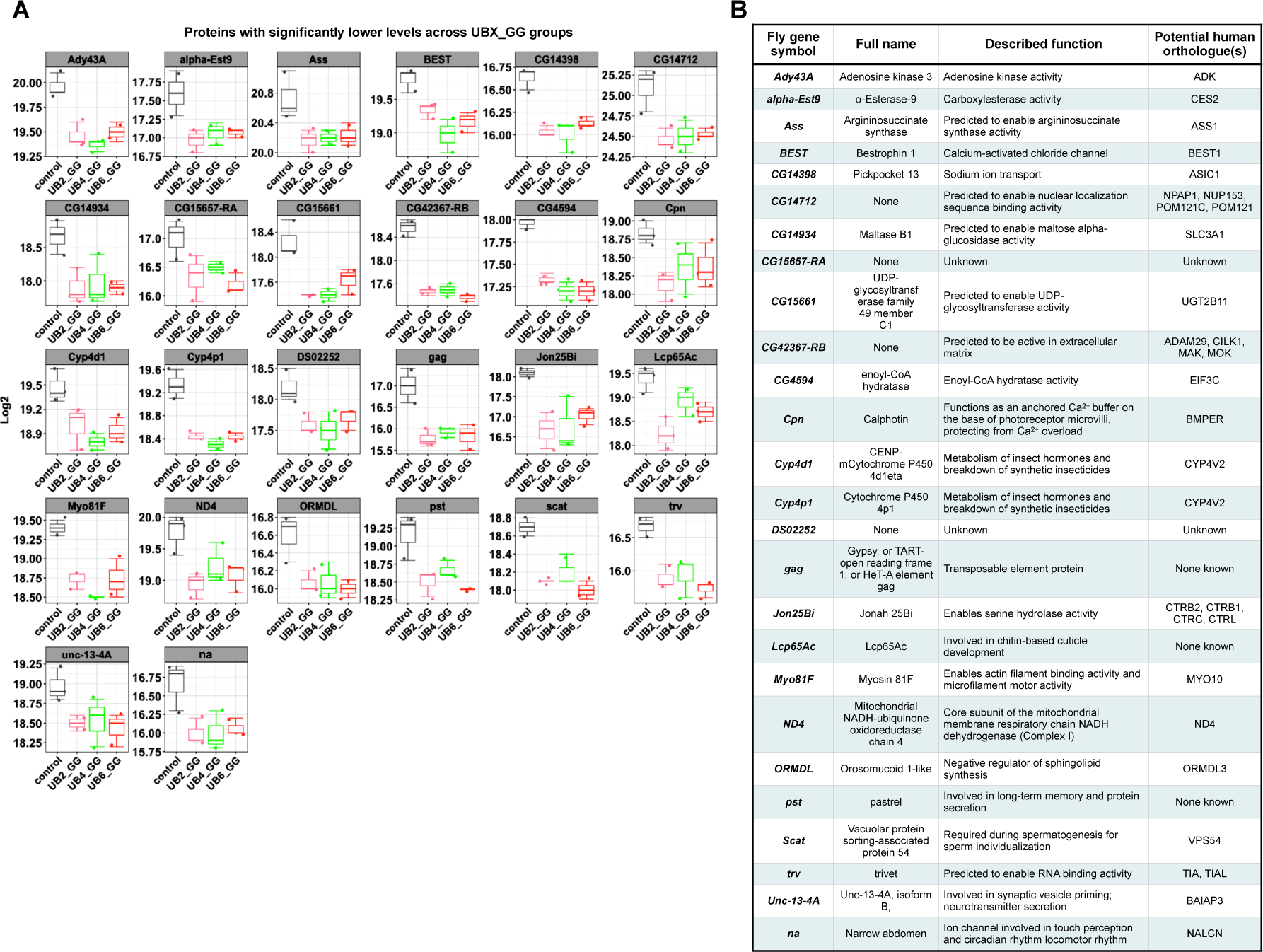
Boxplot of log2 differentially abundant protein expression (A; means -/+ SEM) and their known orthologues and functions (B) whose levels were consistently lower in the presence of UBX_GG. Gene name symbol, described function, and potential human orthologues were sourced from (36). For panel A, individual data points shown represent biological replicates from N=3.

## DISCUSSION

Ubiquitination is a fundamental post-translational modification that controls the function and levels of target proteins. Recently, linear ubiquitin (UB) chains released from ubiquitinated proteins have emerged as an important form of ubiquitin assembly that may play regulatory roles for many cellular processes, such as immune responses and proteostasis (11, 15, 17, 20-24, 44). Our previous studies in *Drosophila melanogaster* demonstrated the versatility of transgene-expressed linear poly-ubiquitin chains that cannot be dismantled into single ubiquitin moieties, which can be conjugated to target proteins or, alternatively, can be themselves ubiquitinated and degraded (20, 28, 29, 45).

Here, we set to determine what protein substrates are modulated by linear UB chains. To this purpose, we examined the changes in protein levels induced by non-cleavable, linear UB chains comprising 2, 4 or 6 ubiquitin moieties when expressed ubiquitously in *Drosophila*. We found numerous proteins – and corresponding processes and pathways – whose levels were changed in the same or opposite direction by the linear UB chains with distinct length and capacity for ligation onto target proteins. Collectively, we found similarities in the proteins and associated cellular functions in the presence of linear UB chains lacking a terminal “GG” *versus* the linear UB chains with “GG”. Differences also emerged in the proteins and cellular processes impacted by linear chains with terminal “GG” motifs but consisting of different UB lengths. We interpret these results to indicate that linear UB species of distinct lengths have specific roles *in vivo* in *Drosophila*, and that they have the potential to modulate a wide range of cellular activities.

To obtain enough material for TMT mass spectrometry, we utilized a ubiquitous Gal4 to drive the transgenic expression of linear poly-UB in *Drosophila* throughout development and in adulthood. While this analysis provides insight into the proteome subsets that are generally and consistently regulated by linear UB chains across multiple tissue, it is possible that the modality of their regulation may differ in specific tissues. Consequently, it is likely that many more proteins could be regulated by linear UB chains in specific tissues and that such tissue-specific targets might be missed by the current analysis of organism-wide changes. Likewise, the proteins modulated by linear UB chains may also change depending on environmental stimuli (e.g., diets and infections) and settings (e.g., aging versus development). Nonetheless, despite the limitations of the current analysis, this study provides fundamental insight into the proteins that are generally modulated by linear UB chains and thus enhances our understanding of how this component of the ubiquitin-proteasome systems contributes to maintain cellular homeostasis.

We started this study to further enhance our understanding of the flexibility of the ubiquitin system in vivo. We had originally hypothesized that expression of linear UB that cannot be dismantled into its constituent moieties is toxic to organisms (27). We found that this was not the case, unless the ability of the cell to manipulate the linear chain was severely restricted – meaning that it would not be able to cleave it, conjugate it to another protein, or conjugate ubiquitin onto it (27, 29). We also found that exogenous, linear UB that could not be disassembled, but was able to be conjugated onto other proteins, could indeed do so *in vivo* (27). Because these data challenged the original hypothesis that linear, free UB chains would be toxic, we next wondered how the cell might employ linear UB chains of different lengths, and how these chains may influence cellular processes and homeostasis. As indicated by the quantitative proteomics analyses that we report in this study, it appears that *Drosophila* cells can utilize distinct linear UB chains to modulate many important cellular processes, thus providing insight into the possible roles that endogenous linear UB chains may play in vivo.

In conclusion, we presented evidence that linear ubiquitin chains remodel the proteome and influence the levels of hundreds of important regulators of homeostasis in *Drosophila*.

## METHODS

### Antibodies

Anti-HA (rabbit monoclonal C29F4, 1:500-1000; Cell Signaling Technology); anti-HIS (rabbit monoclonal, D3I1O; 1:500, Cell Signaling Technology); anti-ubiquitin (mouse monoclonal A5, 1:1000, Santa Cruz Biotech); anti-K48 ubiquitin (rabbit monoclonal 8081, 1:500-1000; Cell Signaling Technology); anti-K63 ubiquitin (rabbit monoclonal DA7011, 1:500-1000; Cell Signaling Technology). HRP-conjugated secondary antibodies were used at 1:5000 and were from Jackson ImmunoResearch.

### *Drosophila*-related methods

Flies were mated, reared and maintained in diurnally controlled environments at 25°C. We used cornmeal-based fly media. The transgenic UB6 and UB6_GG lines were previously reported(27, 29). The UB2, UB2_GG, and UB4_GG lines were generated similarly, by sub-cloning the intended linear UB construct into pWalium10.moe. The precise amino acid sequence used is shown for UB2 in supplemental figure 1. The same sequence was duplicated to generate UB4_GG. Transgenic flies were generated through pHIC-31-depndent integration of pWalium10.moe into site attP40 on chromosome 2 of the fly. Upon transformation and confirmation of the insertion and its sequence via PCR and sequencing, all lines were transferred onto the w1118 background. The sqh-Gal4 line was a gift of Dr. Daniel Kiehart (Duke University).

### Western blotting

Five male flies per group were homogenized in hot SDS lysis buffer (50 mM Tris pH 6.8, 2% SDS, 10% glycerol, 100 mM dithiothreitol (DTT)), sonicated, boiled for 10 minutes, and then centrifuged for 10 minutes at 13,300 rpm at room temperature. Samples were electrophoresed using Tris/Glycine gels (Bio-Rad). Western blot imaging was conducted using ChemiDoc (Bio-Rad). For loading controls, we used direct blue stains of total protein by saturating PVDF membranes for 10 min in 0.008% Direct Blue 71 (Sigma-Aldrich) dissolved in 40% ethanol and 10% acetic acid, rinsed with a solution of 40% ethanol/10% acetic acid, then ultra-pure water, and then air dried and imaged.

### Immunoprecipitation

For panels 1D and 1G, ten adult males were homogenized in 500 µL of NETN lysis buffer (50 mM Tris, pH 7.5, 150 mM NaCl, 0.5% Nonidet P-40) supplemented with protease inhibitors (PI without EDTA; Sigma-Aldrich). Homogenates then centrifuged for 10 minutes at 10,000 x g at 4°C. The supernatant was subsequently combined with bead-bound anti-HA antibody for all species, except UB6_GG, which was incubated with Ni-NTA beads (figure 1B; Sigma-Aldrich) and tumbled at 4°C for 4 hours. Beads were rinsed 5 times each with lysis buffer. Bead-bound complexes were eluted through Laemmli buffer (Bio-Rad) and heating at 95°C for 5 minutes. The supernatant was supplemented with 6% SDS to a final concentration of 1.5% SDS and loaded for Western blotting. For immunoprecipitation data shown in figure 1E, the recombinant protein (GST alone or GST-ZnF) was immobilized on glutathione-sepharose beads using protocols we have described in the past (46), and then was supplemented with *Drosophila* lysates (10 whole flies per sample) or whole mouse brain lysates homogenized in NETN + PI at a concentration of 1 mg/mL. Incubation was conducted at 4°C, tumbling for 4 hours. Beads were then rinsed 5X with the lysis buffer and complexes were eluted through Laemmli buffer (Bio-Rad) and heating at 95°C for 5 minutes.

### Protein sample preparation, protein digestion, and peptide isobaric labeling by tandem mass tags

For each TMT *Drosophila* sample, 20 male flies at 10 days of age were collected and homogenized in 8M urea lysis buffer (50 mM HEPES, pH 8.5, 8 M urea) (44, 47). After homogenization with zirconium beads in a NextAdvance bullet blender, 0.5% sodium deoxycholate was added to the tissue homogenates, which were then briefly pelleted to remove cuticle debris. The protein concentration of the lysates was determined by Coomassie-stained short gels using bovine serum albumin (BSA) as standard, as previously described (48). For TMT mass spectrometry, 100ug of each sample was digested with LysC (Wako) at an enzyme-to-substrate ratio of 1:100 (w/w) for 2 hours in the presence of 1 mM DTT. Following this, the samples were diluted to a final 2 M Urea concentration with 50 mM HEPES (pH 8.5), and further digested with trypsin (Promega) at an enzyme-to-substrate ratio of 1:50 (w/w) for at least 3 hours. The peptides were reduced by adding 1 mM DTT for 30 min at room temperature (RT) followed by alkylation with 10 mM iodoacetamide (IAA) for 30 minutes in the dark at RT. The unreacted IAA was quenched with 30 mM DTT for 30 minutes. Finally, the digestion was terminated and acidified by adding trifluoroacetic acid (TFA) to 1%, desalted using C18 cartridges (Harvard Apparatus), and dried by speed vac. The purified peptides were resuspended in 50 mM HEPES (pH 8.5) and labeled with 18-plex Tandem Mass Tag (TMTpro) reagents (ThermoScientific) following the manufacturer’s recommendations and our optimized protocol (49).

### Two-dimensional HPLC and mass spectrometry

The TMT-labeled samples were mixed equally, desalted, and fractionated on an offline HPLC (Agilent 1220) by using basic pH reverse-phase liquid chromatography (pH 8.0, XBridge C18 column, 4.6 mm × 25 cm, 3.5 μm particle size, Waters). The fractions were dried and resuspended in 5% formic acid and analyzed by acidic pH reverse phase LC-MS/MS analysis. The peptide samples were loaded on a nanoscale capillary reverse phase C18 column (Thermo PepMap RSLC column, 75 um ID × 15 cm, 2 μm C18 resin) by an HPLC system (Thermo Ultimate 3000) and eluted by a 100-min gradient. The eluted peptides were ionized by electrospray ionization and detected by an inline Q-Exactive HF mass spectrometer (ThermoScientific). The mass spectrometer was operated in data-dependent mode with a survey scan in Orbitrap (60,000 resolution, 1 × 10^6^ AGC target and 50 ms maximal ion time) and MS/MS high-resolution scans (60,000 resolution, 1 × 10^5^ AGC target, 110 ms maximal ion time, 32 HCD normalized collision energy, 1 *m*/*z* isolation window, and 10 s dynamic exclusion).

### MS data analyses

The MS/MS raw files were processed by the tag-based hybrid search engine, JUMP (50). The raw data were searched against the UniProt *Drosophila* databases concatenated with a reversed decoy database for evaluating false discovery rates. Searches were performed by using a 25-ppm mass tolerance for both precursor and product ions, fully tryptic restriction with two maximal missed cleavages, three maximal modification sites, and the assignment of *a*, *b*, and *y* ions. TMT tags on Lys and N-termini (+304.20715 Da) were used for static modifications and Met oxidation (+15.99492 Da) was considered as a dynamic modification. Matched MS/MS spectra were filtered by mass accuracy and matching scores to reduce protein false discovery rate to ∼1%. Proteins were quantified by summing reporter ion intensities across all matched PSMs using the JUMP software suite (51).

### Supervised and over-representation analysis

Raw values from MS data analyses were used to run a PCA using the ‘prcomp’ function in R software. Log2FC values of the UBX group in comparison to control was used to plot a heatmap (ComplexHeatmaps in R) (55, 57) using protein IDs curated from flybase.org and annotated to relevant pathways addressed in the text. Differentially abundant proteins were identified as proteins that were significantly different from control (p<0.05) with log2FC >/= 0.5. GeneOverlap package (58) was used visualize protein ID overlaps and to calculate significance using a genome size background of 8000 genes. Differentially abundant genes were used to determine pathway enrichment by employing GeneOntology (59, 62) Kyoto Encyclopedia of Genes and Genomes (KEGG) (63, 65) and the clusterprofiler (66, 67) packages in R.

## ACKNOWLEDGEMENTS

We thank Drs. Jessica R. Blount and Kozeta Libohova (Wayne State University) for their assistance with *Drosophila* sample collection.

## FUNDING

This work was funded in part by R21NS123147 and R01NS086778 to SVT. The content is solely the responsibility of the authors and does not necessarily represent the official views of the National Institutes of Health. Research at St. Jude Children’s Research Hospital is supported by the ALSAC.

## FIGURE LEGENDS

**Supplemental figure 1:**
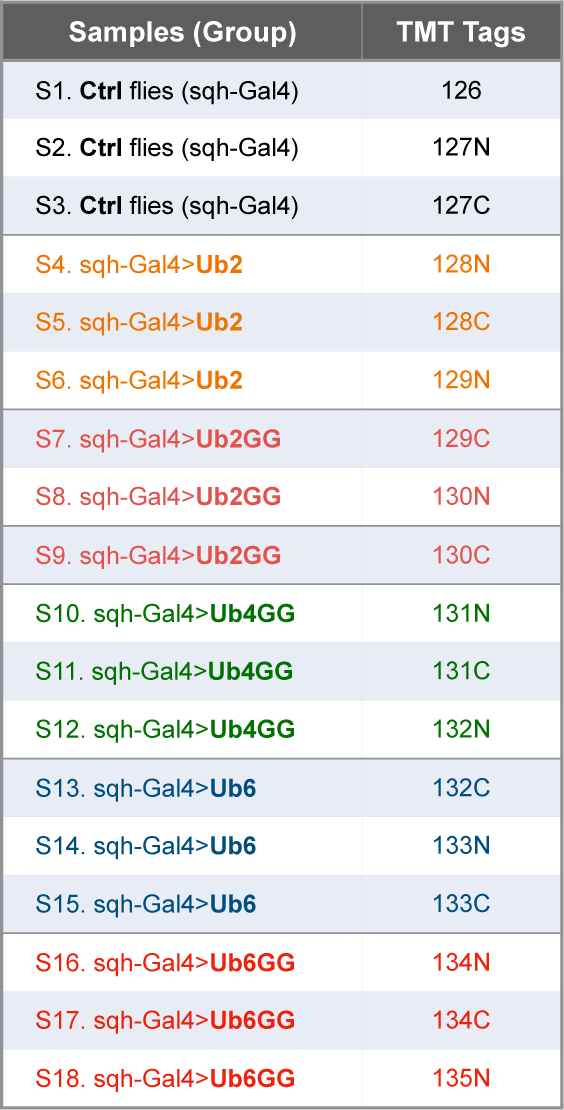
TMT tags used for each group.

**Supplemental figure 2:**
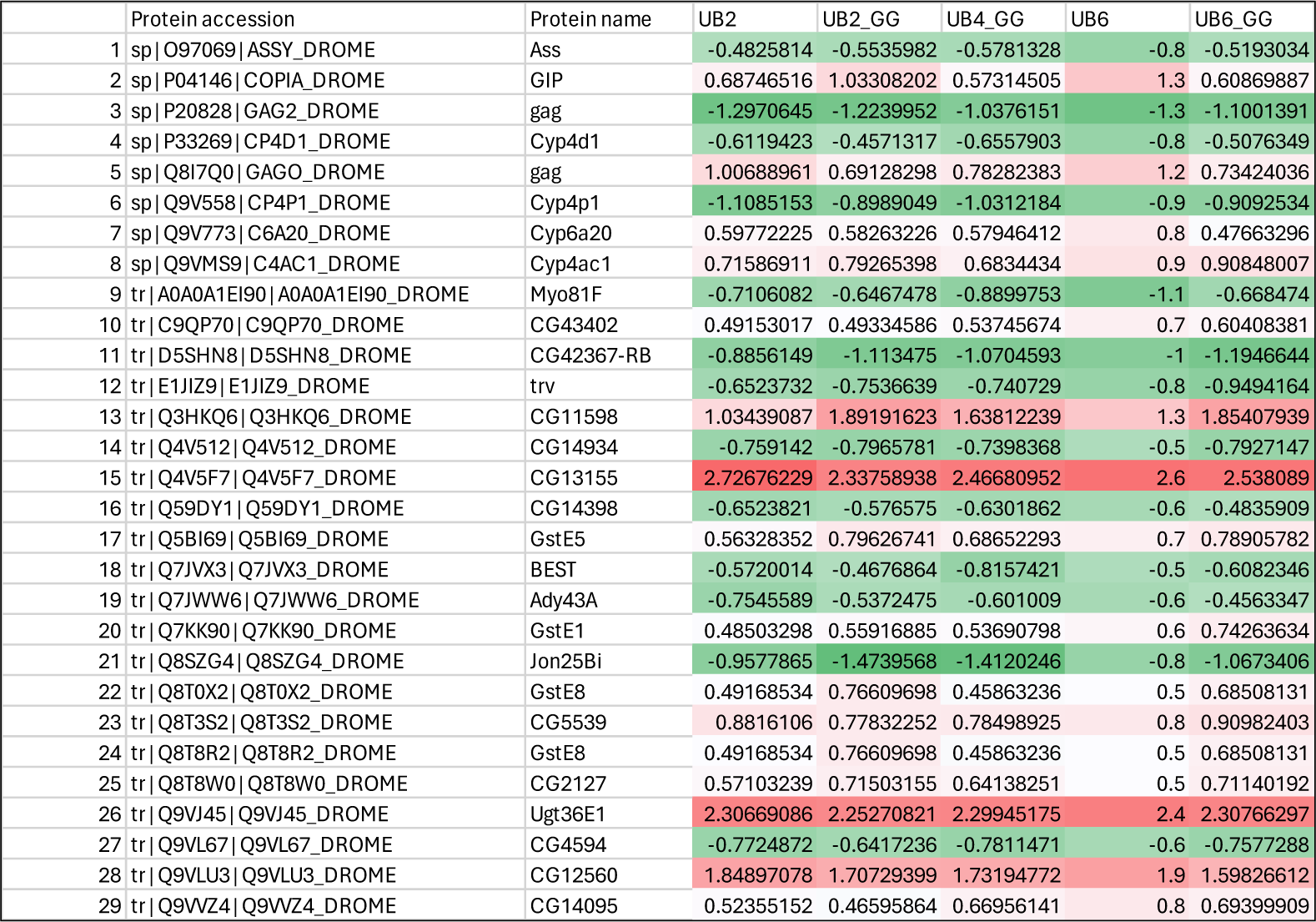
List of proteins intersecting in figure 2A and their directionality of change.

**Supplemental figure 3:**
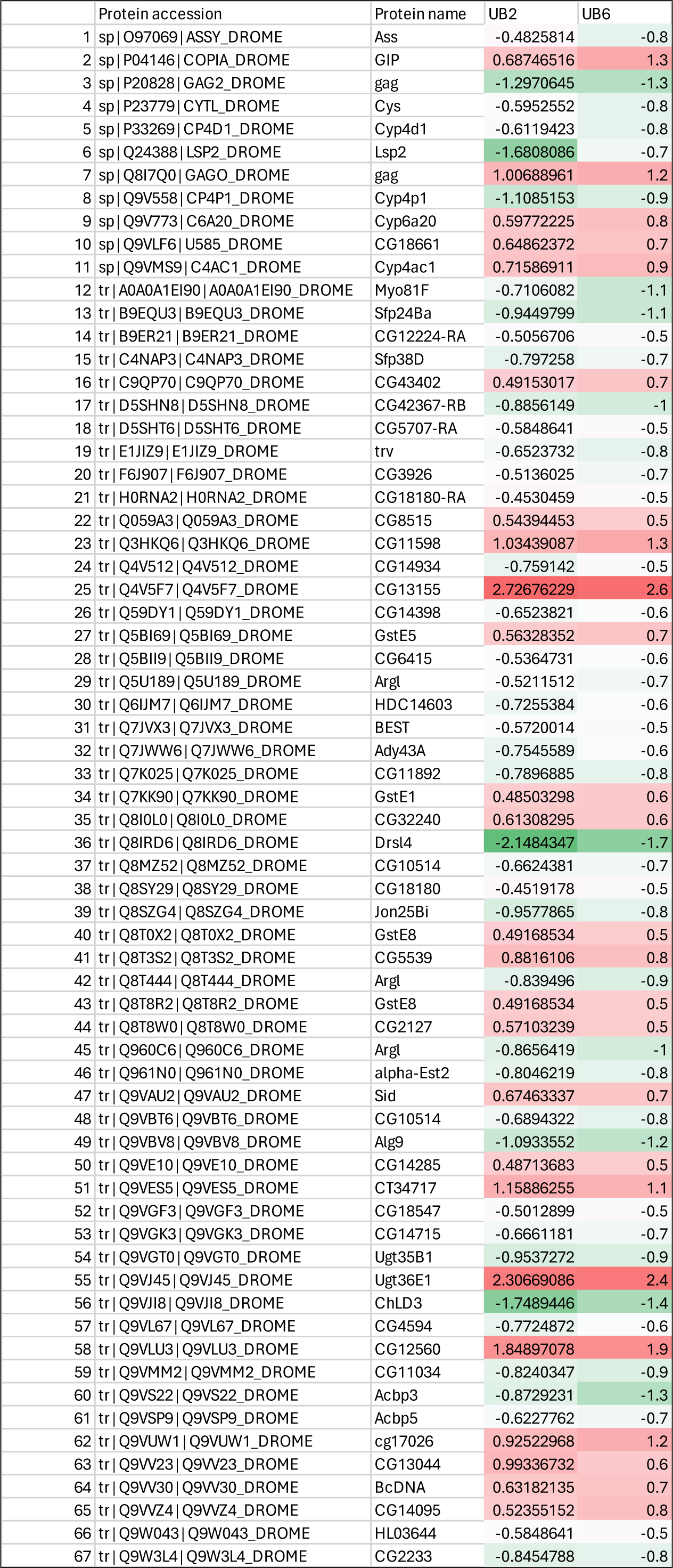
List of proteins intersecting in figure 2B and their directionality of change.

**Supplemental figure 4:**
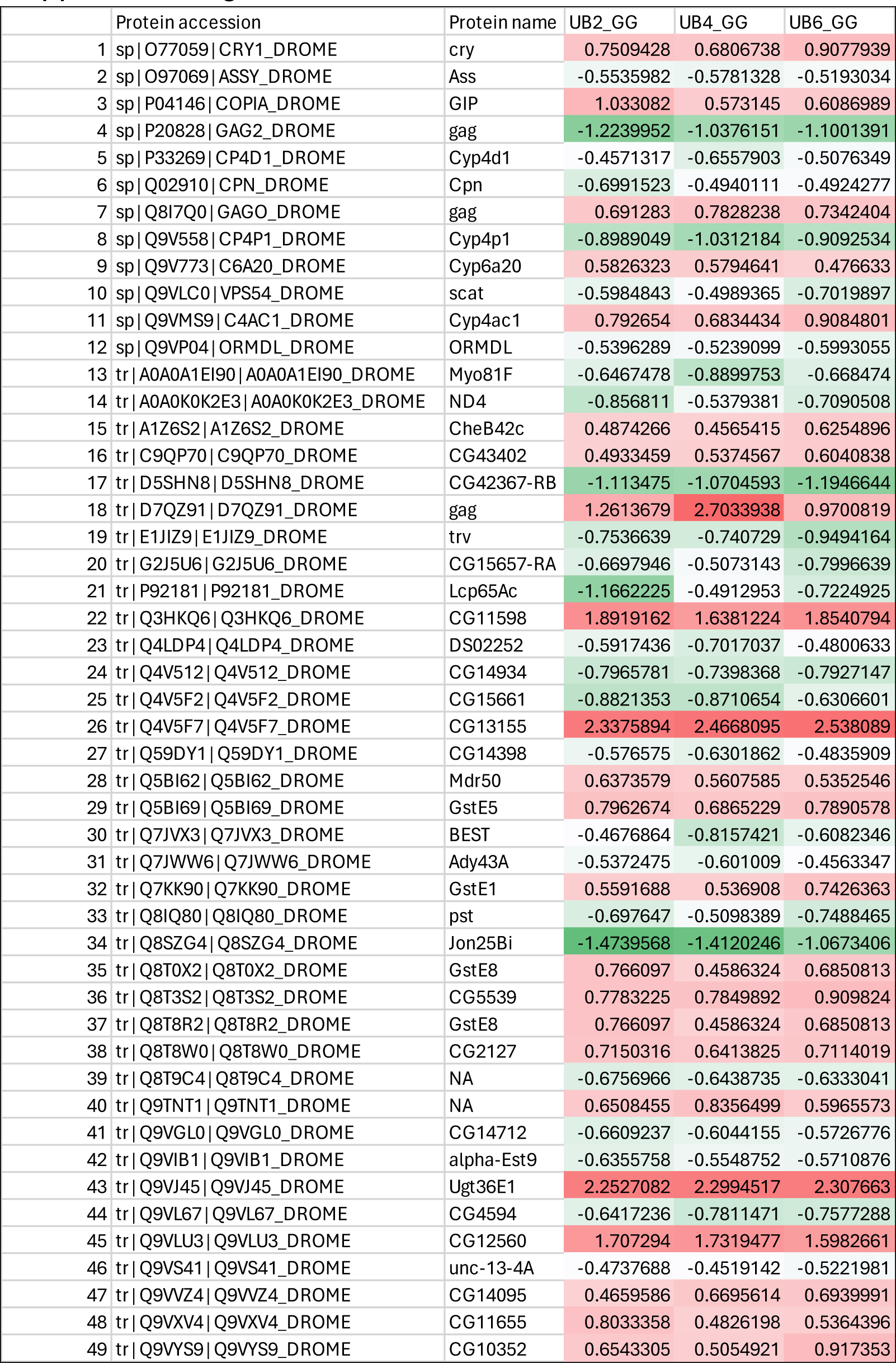
List of proteins intersecting in figure 2C and their directionality of change.

**Supplemental figure 5:**
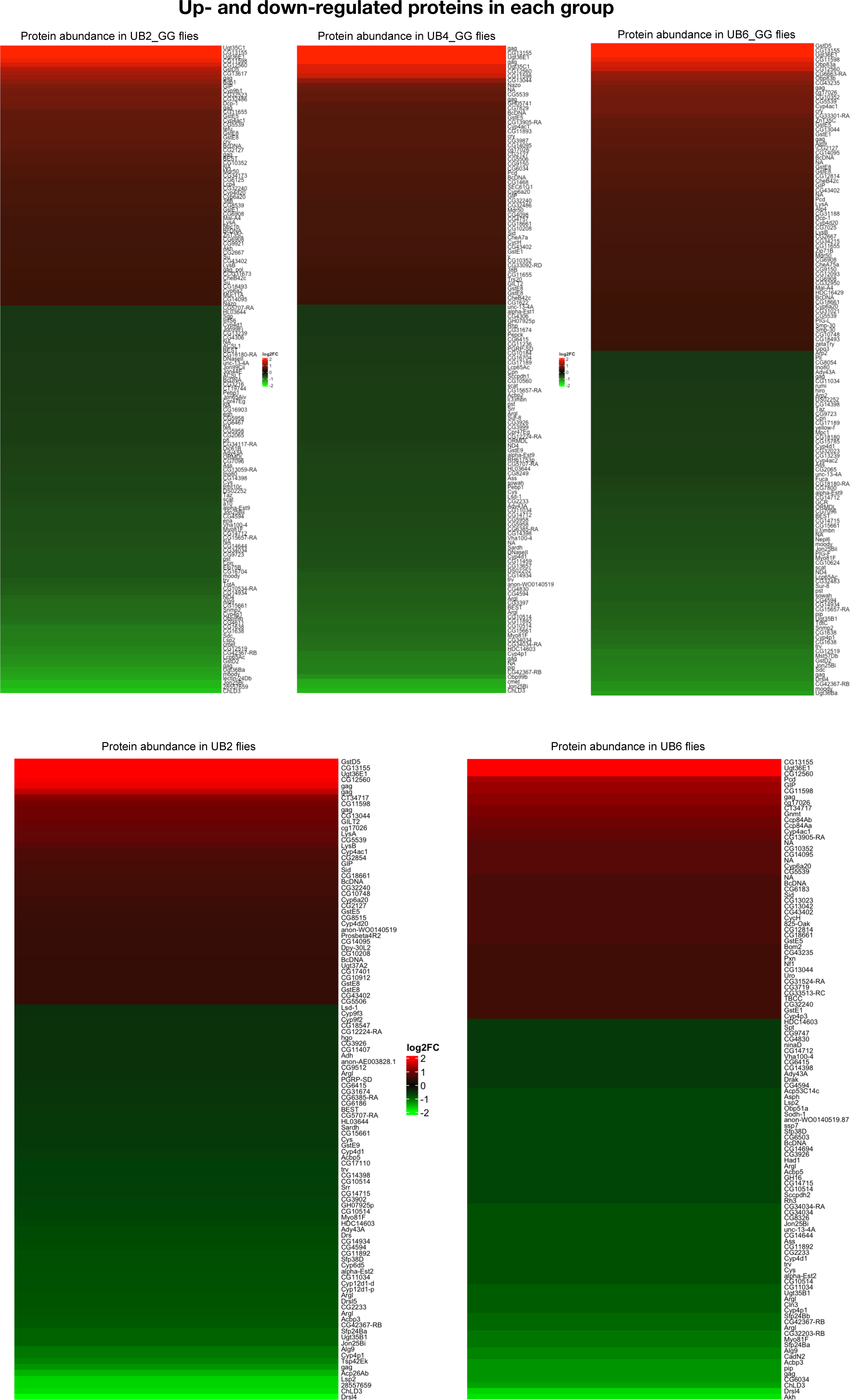
Heatmaps of all proteins that were up- or down-regulated in the presence of specific UBX linear proteins.

## Notes

### Competing Interest Statement

The authors have declared no competing interest.

